# DECODE: DEcomposition of COupled and DEcoupled Brain States: Separating Structure–Coupled and Structure–Decoupled Functional Organization in Alzheimer’s Disease

**DOI:** 10.64898/2026.11.09.765343

**Authors:** Mahshid Fouladivanda, Armin Iraji, Souvik Phadikar, Ram Balm, Bikesh Bimali, Tulay Adali, Vince D. Calhoun

**Affiliations:** Tri-Institutional Center for Translational Research in Neuroimaging and Data Science (Georgia State University, Georgia Institute of Technology, Emory University), Atlanta, USA; Department of Computer Science, Georgia State University, Atlanta, GA, USA; School of Electrical and Computer Engineering, Georgia Institute of Technology, Atlanta, USA; Department of Computer Science and Electrical Engineering, University of Maryland Baltimore County, Baltimore, MD, USA

**Keywords:** Latent-space decomposition, Functional states, Structure-function, Structure-Coupled, Structure-Decoupled

## Abstract

Brain functional activity emerges within an underlying structural scaffold. While structural network connectivity provides a relatively stable anatomical backbone for large-scale communication, functional interactions are high-dimensional and temporally dynamic, forming recurrent patterns that can be represented in a latent space of transient functional states. Yet, how these transient functional states are organized and the extent to which they remain constrained by structural network connectivity or become relatively decoupled from structural organization over time remain poorly understood. To address this gap, we introduce DECODE (DEcomposition of COupled and DEcoupled Brain States), a data-driven latent-space framework that uses independent component analysis (ICA) and multimodal data to decompose resting-state dynamic functional network connectivity into a lower-dimensional representation of transient functional states. These states capture temporally varying structure-coupled and structure-decoupled patterns using diffusion MRI-derived structural network connectivity priors. Structure-coupled states reflect functional configurations more strongly anchored to major structural motifs, whereas structure-decoupled states represent configurations less constrained by the identified structural organization, while potentially remaining within structurally defined dynamical bounds. DECODE identifies reproducible structure-coupled and structure-decoupled states, with mean squared error reconstruction of dynamic information < 0.1 and adjusted R^2^ = 0.4. Applied to Alzheimer’s disease (AD), DECODE revealed stronger group differences (FDR-corrected p-value 0.05) in both connectivity organization and temporal dynamics for decoupled states. These findings suggest that functional states less constrained by identified structural organization may be more vulnerable to AD-related disruption, whereas structure-coupled states may be relatively preserved. More broadly, DECODE provides a reproducible and generalizable framework for quantifying how functional dynamics are shaped by, or diverge from, structural organization by incorporating a latent representation of the structural connectome as an informative prior. Our findings established structural and functional alignment variation by distinguishing dynamic functional patterns closely coupled with structure from those that deviate from it.

## Introduction

Structural and functional network connectivity offer complementary views of brain organization that is not well understood how they are aligned. Structural network connectivity reflects the anatomical scaffold supporting communication between brain regions, whereas functional network connectivity captures coordinated neural activity across them [1, 2]. Although related, they are not equivalent: structural network connectivity is a relatively fixed pattern, whereas functional network connectivity is a temporal synchronization activity between brain regions [3, 4]. Rather than a single configuration, functional network connectivity exhibits recurring patterns of whole-brain organization, commonly characterized using dynamic functional network connectivity (dFNC) [1-3]. These recurring patterns define a lower-dimensional latent space of functional organization, representing distinct modes of coordinated brain activity. An important question is how structural network connectivity shapes this latent space functional organization. Anatomical structure constrains functional interactions without fully determining them. Directly connected regions often exhibit stronger functional network connectivity, while indirect or polysynaptic pathways can support a broader range of functional relationships [5]. Thus, functional organization may be constrained by structural architecture to varying degrees rather than determined by direct anatomical correspondence.

Previous studies have often assessed structure-function associations using regional spatial correlations or related statistical measures [5-7], compared structural and functional modalities after separating unimodal analyses [2, 8], or examined their relationship through alternative formulation such as harmonic analysis [9]. More recent graph-based approaches, including the structural decoupling index (SDI), decompose functional signals according to their alignment with structural graph organization [10]. Together, these approaches demonstrate that functional organization can be systematically related to structural architecture, but primarily quantify the strength or degree of structure-function association rather than how structural constraints are distributed across different functional configurations [3]. Consequently, direct structure-function coupling estimates can be potentially affecting the interpretability and generalizability of the measurements. This concern highlights an important limitation of methods that rely on direct region-wise correspondence between structural and functional network connectivity profiles [11, 12]. In addition, they typically summarize this relationship using static or regional descriptors and may not fully capture its dynamic, state-dependent nature. Even when structure-function coupling is examined using alternative formulations, most approaches still quantify association rather than decompose functional organization into components with distinct degrees of structural alignment.

This limitation is particularly relevant for studies of dynamic functional network connectivity. Recurrent brain states are typically estimated from functional data alone, without explicit structural priors [13-15], while multimodal studies often stop at cross-modal comparison rather than joint decomposition. As a result, it remains unclear how much of recurrent functional brain-state organization is directly constrained by anatomical structure and how much reflects more flexible, relatively decoupled functional processes [1, 2]. A dynamic multimodal decomposition model that can jointly identify structure-coupled and structure-decoupled brain states may therefore provide a more mechanistic account of how brain dynamics transition from structural constraint to functional flexibility [2, 10].

To address this gap, we introduce DECODE (Decomposition of Coupled and Decoupled Brain States), a data-driven multimodal framework that estimates a latent space of dynamic brain states and decomposes it according to the structural constraint. DECODE first estimates structural connectivity patterns capturing coordinated anatomical organization, which are then incorporated as priors within an independent component analysis framework [16-18] to identify functional states that are more strongly anchored to, or relatively less constrained by, structural organization. Importantly, structure-coupled states do not imply one-to-one SC-FC correspondence; rather, they reflect functional configurations aligned with the distributed organization of the structural connectome, potentially through direct and indirect anatomical influences. Thus, DECODE decomposes the latent space of dynamic functional organization into structure-coupled and structure-decoupled states.

We demonstrate the application of DECODE to Alzheimer’s disease (AD), a neurodegenerative disorder characterized by widespread alterations in structural and functional brain organization and cognitive impairment. We use DECODE to quantify dysfunction in structure-coupled and structure-decoupled brain states in AD. Although demonstrated in AD, DECODE is not disease-specific and provides a general framework for investigating how dynamic functional organization is shaped by, or diverges from, structural network connectivity across populations and brain disorders.

## Materials and Methods

### Participants

We obtained the data from the Alzheimer’s Disease Neuroimaging Initiative (ADNI) database (https://adni.loni.usc.edu/), including Cognitively Normal (CN), Mild Cognitive Impairment (MCI), and Alzheimer’s disease (AD) individuals.

### Preprocessing

Resting-state fMRI data were preprocessed using a standard pipeline similar to [19], including motion and distortion correction with FSL (https://fsl.fmrib.ox.ac.uk/fsl/fslwiki) toolbox, slice timing correction with SPM (https://www.fil.ion.ucl.ac.uk/spm/software/spm12/), spatial normalization to the standard Montreal Neurological Institute (MNI) space, and spatial smoothing using a 6-mm Full-width Half Maximum (FWHM) Gaussian kernel. For the diffusion MRI data we used the pipeline described in [20], to perform skull stripping, eddy current correction with FSL [21], denoising using local principal component analysis (PCA) in DIPY, and nonlinear registration to MNI space to correct for EPI-induced currents.

After preprocessing the data, 2,290 resting-state fMRI sessions from 1,038 unique participants and 2500 diffusion MRI sessions from 1,097 unique participants remained. fMRI data distribution over the three diagnostic groups are 1044 CN sessions, 734 sessions MCI, and 512 sessions with AD, more details in demographic information of the fMRI data, including age and sex, fMRI data is summarized in Table 1.

**Table 1.**
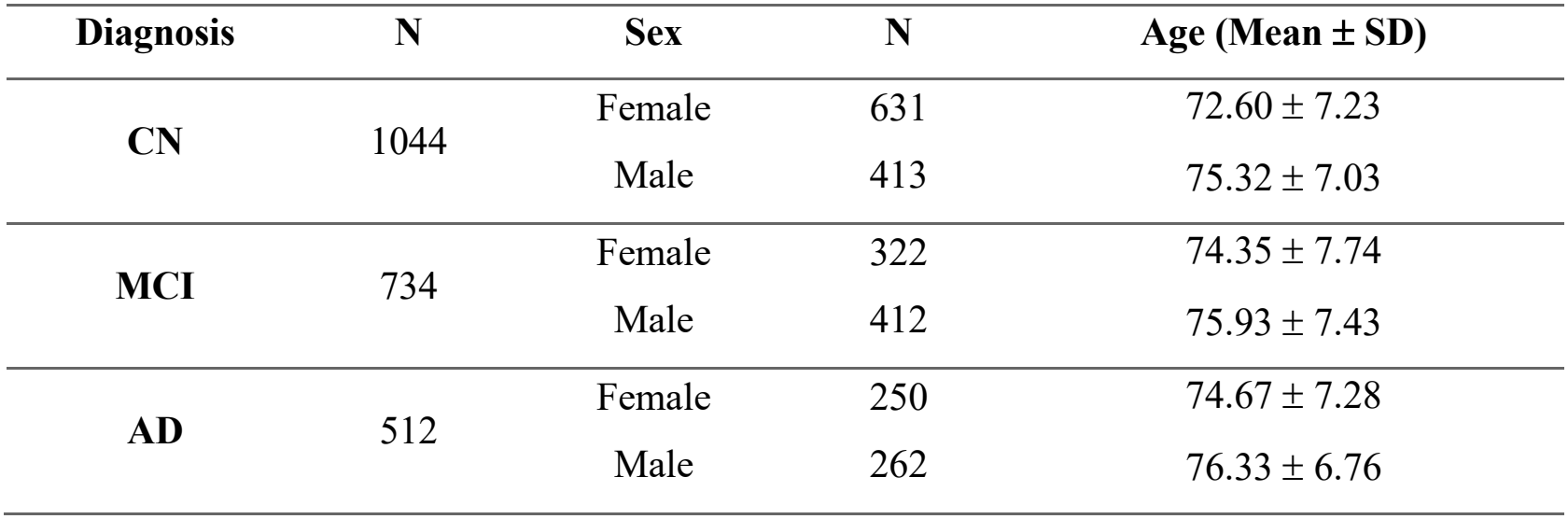
Demographic information of the fMRI data.

### Dynamic functional network connectivity

Each preprocessed fMRI scan was analyzed using the NeuroMark pipeline [18], a unified independent components analysis (ICA)-based framework for estimating intrinsic connectivity networks (ICNs). We used NeuroMark_fMRI_1.0 as the template to estimate 53 ICNs organized into seven canonical functional domains: SC (Subcortical), AUD (auditory), SM (Sensorimotor), VS (Visual), CC (Cognitive Control), DMN (default-mode network), and CB (Cerebellum). ICNs represent spatially distributed networks with coherent temporal activity, each characterized by a spatial map and am associated time course reflecting its temporal activity (TA).

We then estimated the dynamic functional network connectivity (dFNC) using a sliding-window approach. A rectangular window with length of 30 repetition time (TR = 1.5 seconds) was tapered by convolution with a Gaussian kernel (σ = 3 TRs) [2, 22] and slid across each TA in steps of 1 TR, segmenting the time course of each ICNs into short intervals (windowed TA). Pairwise Pearson correlations were computed within each window, yielding a 53 × 53 × 176 dFNC matrix per individual.

### Structural network connectivity

Structural network connectivity (SNC) matrices were constructed using a whole-brain fiber tractography on the preprocessed dMRI images. First, we applied diffusion tensor fitting, implemented in dtifit under FSL toolbox [21], to estimate voxel-wise tensors. We then performed a deterministic tractography algorithm in MRtrix3 (https://www.mrtrix.org/) to generate whole-brain fiber tracts in native space, using a step size of 0.5 mm, an anisotropy threshold of 0.2, and an angular threshold of 60°. To ensure correspondence with the networks used for dFNC estimation, the normalized NeuroMark_fMRI_1.0 template was inverse transformed into each individual’s native space, yielding spatial maps of the 53 ICNs [18, 23]. For each pair of ICNs, the number of fiber tracts connecting the two ICNs was computed, constructing an individual-level 53 × 53 structural connectivity matrix.

### Structural latent space estimation

To derive anatomically informed priors for DECODE, each SNC matrix was vectorized by extracting its upper triangle (1,378 unique connections), and these vectors were concatenated across individuals to form a group-level matrix (N×1378). We then applied group-level independent component analysis (gICA) [16], implemented with the Infomax algorithm [24, 25] in the GIFT toolbox (https://github.com/trendscenter/gift), to estimate a set of structural latent components. Structural latent components (SLC_k_) are ICA-derived components representing statistically independent structural connectivity patterns shared across individuals. Each SLC_k_ represents a latent space of the *k*^th^ covarying structural pattern and together they provide a compact representation of the large-scale anatomical scaffold. Components stability was assessed using ICASSO [16, 26] with 100 bootstrap iterations. The optimal number (*k*) of the latent-space components was determined using the elbow criterion implemented in GIFT, yielding five structural latent components in the present study.

Rather than using anatomy only as a post hoc reference, applying post hoc correlation, or late fusion, DECODE uses this group-derived latent components as priors to constrain the decomposition of dynamic functional organization. Because the prior is defined on latent components rather than on individual SNC matrices, and enters the decomposition adaptively rather than through fixed regression, the model does not impose a direct one-to-one correspondence between structural and functional connectivity.

### Structural – functional states decomposition

#### Model formulation

The core contribution of DECODE is the decomposition of dynamic functional network connectivity (dFNC) into components that coupled with structural organization and components that do not. We assume that dynamic brain activity can be expressed as a linear mixture of two sets of connectivity patterns: those coupled to the structural scaffold, captured by the group-level structural latent components, and those not explained by this structural prior. We refer to these as structure-coupled and structure-decoupled states, respectively. Main pipeline of the proposed model is displayed in Figure 1.

**Figure 1.**
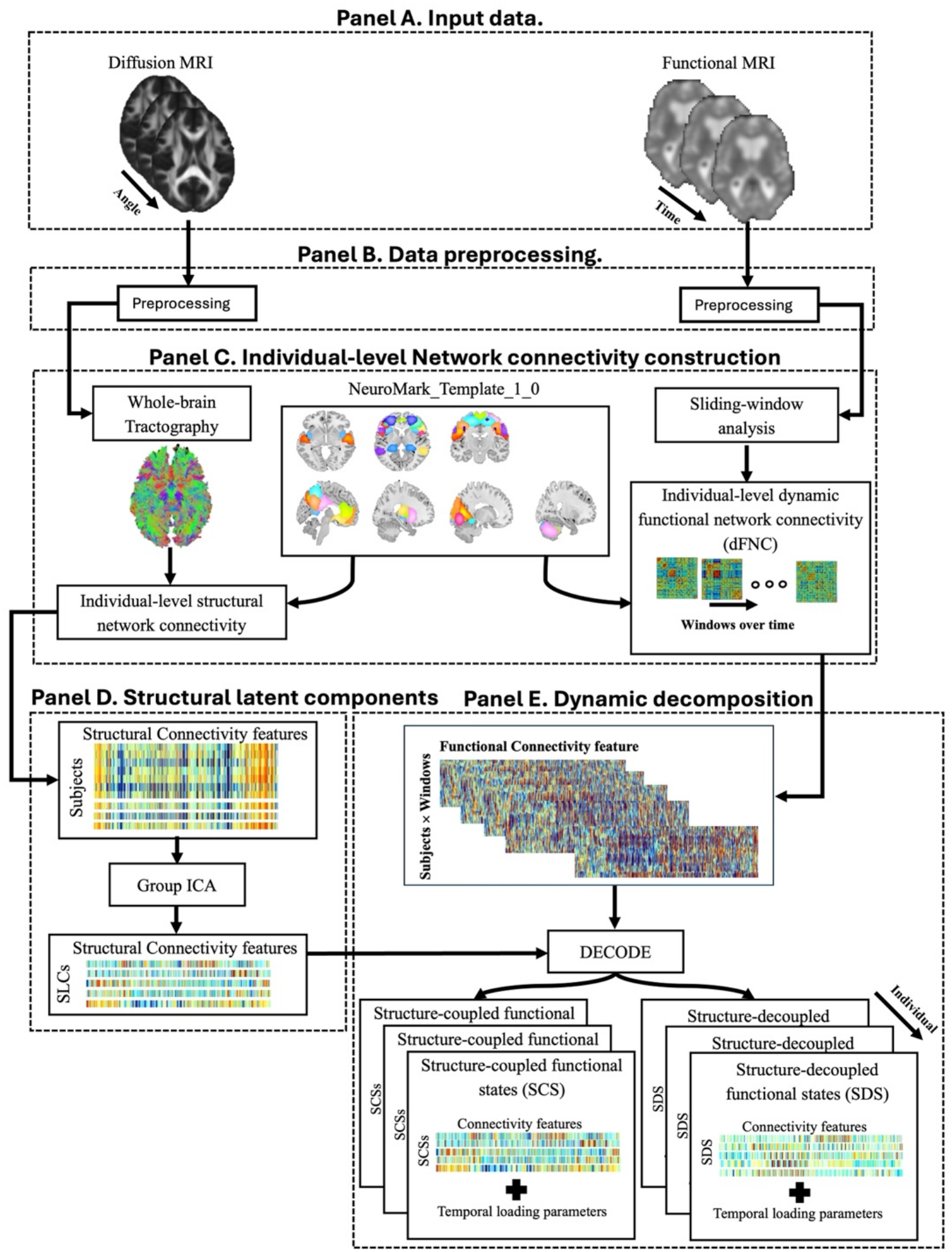
The overall framework for the proposed DECODE model is displayed. Panel A illustrates the input data including diffusion MRI and functional MRI images. Each data was preprocessed using separate pipelines as is shown in Panel B. Next, network connectivity analysis was performed in parallel to estimate individual-level structural network connectivity and dynamic functional network connectivity. As shown in Panel C, combination of whole brain tractography and NeuroMark_fMRI_1_0 template generates individual-level structural network connectivity. Using NeuroMark_fMRI_1_0 template along with sliding window analysis, we defined short-time interval brain activities which a functional network connectivity was estimated per window by computing Pearson’s correlation of paired regions in template. Then, structural latent components (SLCs) were derived from the structural connectivity information to define structural priors that shape brain dynamic communication pathways. These latent components were determined by applying group ICA on individual-level structural network connectivity in Panel D. Finally, in Panel E, estimated dFNC matrices along with the SLCs were guided the decomposition procedure under the DECODE model to uncover two set of reoccurring multimodal latent space states; Structure-Coupled States (SCS) and Structure-Decoupled States (SDS).

Let ***Y*** ∈ ℝ^*T*×*I*^ denote the vectorized dFNC matrices, where *T* = 176, is the number of windows and *I =* 1378 is the number of unique connectivity edges. ***Z*** ∈ ℝ^*K*×*I*^ denotes the ***K*** = [1, 2, … , *k*] group-level structural latent components (SLC), which is fixed. We model the unexplained functional organization by direct structural organization as a second set of latent components, ***X*** ∈ ℝ^*K’*×*I*^ estimated from the dFNCs, where ***K***^*’*^ = [1, 2, … , *k*^*’*^]. DECODE then expresses dFNC as

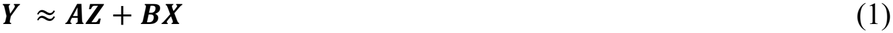

where ***A*** ∈ ℝ^*T*×*K*^ and ***B*** ∈ ℝ^*T*×*K’*^ are mixing matrices, whose rows give the time-resolved loadings of the structure-coupled and structure-decoupled components, respectively.

#### Optimization

We propose an ICA-based model to recover the mixing matrices **A** and **B** from Eq. 1. Previous constrained ICA studies [17, 27-29] have predominantly used unimodal data, most often focusing on functional information alone. We address this limitation with a multimodal decomposition that decomposes the dFNC matrices into latent-space recurring patterns over time, using SLC as prior information. The aim is to find two complementary subspaces through a joint learning procedure, yielding two sets of independent recurring connectivity patterns (states): structure-coupled and structure-decoupled.

The optimization proceeds in three steps:

1. Learn **A** by jointly estimating a structure-function coupled latent space.
2. Compute the unexplained dynamic functional information not reconstructed by the structure-coupled states.
3. Decompose this unconstructed information into a latent space to learn **B**.

Structure-coupled subspace is determined by directly coupling structural and functional connectivity information under a joint optimization process to learn a mixing matrix **A**. By simultaneously maximizing the statistical independence of the estimated components in (2) and their similarity to the structural latent components (SLC), the resulting states preserve alignment with the structural scaffold while remaining data-driven with respect to functional dynamics. However, structure-decoupled space captures the dynamic functional information that is not projected onto the structural latent components. The objective is to identify this unexplained information and transform it into a latent space that captures coherent and recurring patterns.

We define the following three objectives in (2) to assess the decomposition of the dFNC into complementary subspaces.

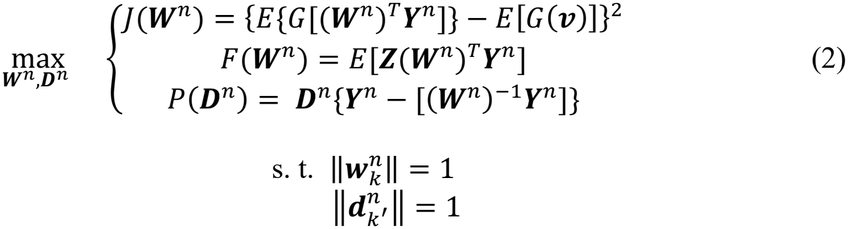

where, ***W***^*n*^ = (***A***^*n*^) ^−1^ and ***D***^*n*^ = (***B***^*n*^)^−1^ denote the estimated unmixing vector for subject *n*. In (2), 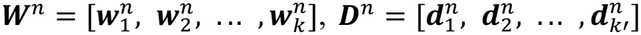 , *k* and *k*^*’*^ index the structure-coupled states and structure-decoupled states, and ***Y***^*n*^ is the whitend dFNC matrix. The number of the structure-coupled states (***K***) corresponds to the number of structural latent components (here, five shown with ***Z***), while the number of structure-decoupled states was determined using the elbow criterion (also five in this study). *E*[.] is the expectation operator, *G* is the nonquadratic contrast function G(**u**) = (1/a_1_) log[cosh(au)], with a_1_ = 1, used to approximate negentropy [30, 31], and ***v*** is a random Gaussian variable with zero mean and unit variance.

*J*(.) promots statistical independence [18] leaning to the dynamic functional information, *F*(.), enforces similarity with structural prior, together they drive the learning of the mixing matrix ***A***, transforming ***Y***^*n*^ into the structure-coupled space using the optimized ***W***^*n*^. *P*(.) operates on the portion of ***Y***^*n*^ that is not accounted for by ***W***^*n*^. The inner product with ***D***^*n*^ is then measures how well this candidate unmixing direction aligns with this unexplained data, serving as a criterion for identifying latent space states that ***W***^*n*^ alone does not capture.

#### Solving the multi-objective problem

To solve the proposed multi-objective problem, we first simplify our problem into the two objectives: one accounting for structure-coupled learning, and one representing the dynamic information not explained by the structure-coupled states.

For the structure-coupled learning, we applied a widely used linear weighted sum of the objectives (3). To prevent bias arising from differing objective scales, *J*(.) and *F*(.) were normalized independently prior to combination, following [17].

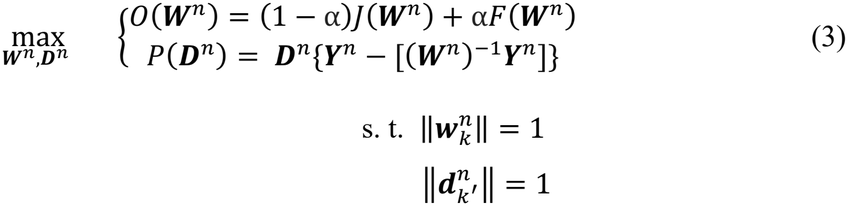

where the mixing parameter (*α* > 0) controls the relative contribution of structural similarity versus statistical independence in the resulting components. In this study, *α* was fixed at 0.5 to provide balanced weighting of the two objectives, preventing either the structural similarity or statistical independence objective from dominating the optimization.

For the initialization of the ***W***^*n*^, a data-driven closed-form solution inspired from [17] was used (4):

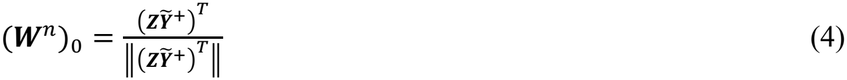

where 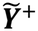 is the pseudo-inverse matrix of the whitened ***Y***^*n*^, such that *F*(***W***^*n*^) ∈ [0,1] at initialization. Also, ***Z*** is z-scored prior to optimization. Optimization was performed using gradient ascent until convergence. Consequently, an optimized unmixing matrix ***W***^*n*^ = (***A***^*n*^)^−1^ was thereby obtained for each dFNC matrix (***Y***^*n*^), indication the contribution of the direct structure-coupled states. Using the estimated unmixing matrix, we separated each dFNC into structure-coupled latent-space states (5) and the unexplained information which is failed to transform to the structural latent components (6). Regressing out the reconstructed structure-coupled states (***C***^*n*^), we captured the unexplained dynamic functional information (***R***^*n*^).

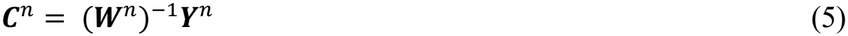

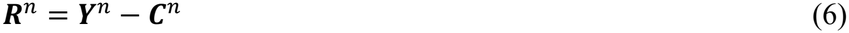

We then estimated a latent space for the unexplained dynamic functional information using *P*(***D***^*n*^) from (3). To solve *P*(***D***^*n*^), we applied a group-level ICA (gICA) model with the Infomax algorithm, yielding the structure-decoupled functional states. Structure-decoupled functional states are a group-level projection of the unexplained functional information into a latent space. Using gICA, we identify a set of statistically independent components representing covarying, co-occurring connectivity patterns across the group. An optimized unmixing matrix, ***D***^*n*^ = (***B***^*n*^)^−1^ was obtained for each dFNC matrix:

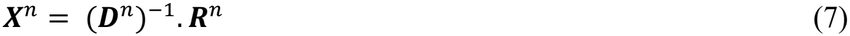

These components represent recurring functional patterns not directly explained by the structural priors. The number of structure-decoupled states was determined using the elbow criterion and was five in this study. Stability was again assessed using ICASSO with 100 bootstrap iterations. To project the group-level structure-decoupled latent space to the individual-level, we applied dual-regression based back-reconstruction model using dual-regression [32].

The proposed DECODE model enables to move beyond conventional coupling analyses by explicitly separating dynamic functional organization into structure-coupled and structure-decoupled latent connectivity pattern (***C***^*n*^ and ***X***^*n*^) and quantifying how each contributes over time through the corresponding mixing matrices (***A***^*n*^ and ***B***^*n*^) at the individual level.

#### Model validation

To assess the generalizability, reproducibility, and robustness of the proposed model, we employed a single hold-out validation strategy combined with the ICASSO approach. Here, generalizability refers to the model’s ability to capture consistent patterns in unseen data, reproducibility reflects the consistency of estimated components across independent data subsets, and robustness refers to the stability of estimated components under variations in data sampling. For the hold-out validation, the dataset was randomly divided into two non-overlapping subsets, with 70% of individuals assigned to the training group and the remaining 30% to the test group.

We first evaluated the generalizability and reproducibility of the structural latent components (SLCs) by estimating them independently for the test and train groups. Group ICA was applied separately to each group using a model order of five, consistent with the SLCs estimated from the full sample. To assess the robustness against sampling variability, we employed the ICASSO approach [16, 26] with 100 bootstrap repetition and captured components with a stability index (Qi) > 0.8. We then quantified the similarity between the SLCs estimated from the test- and train-groups and those derived from the full sample using Pearson’s correlation coefficient.

Similarly, we examined the generalizability, reproducibility and robustness of the structure-coupled and structure-decoupled dynamic states (SCSs and SDSs). Specifically, the dFNC of the test group was decomposed using the SLCs estimated from the train group, and the resulting SCSs and SDSs were compared with their corresponding states derived from the full sample using Pearson’s correlation coefficient.

To evaluate the representational performance of DECODE, we compared explained variance (R^2^) across two decomposition schemes: inter-group and intra-group. In the inter-group scheme, SLCs estimated from the train group were used to decompose the test group dFNC, and vice versa. In the intra-group scheme, each group’s dFNC was decomposed using its own SLCs. Both schemes were benchmarked against two reference approaches: DECODE applied to the full sample and a conventional unimodal functional-state estimation approach based on k-means clustering [13].

To further assess reconstruction accuracy, we quantified the mean squared error (MSE) between the original and reconstructed dFNC. The dFNC of the test group was decomposed into SCSs and SDSs using parameters learned from the train group in (8), and the reconstructed dFNC was compared to the original to compute the MSE as in (9).

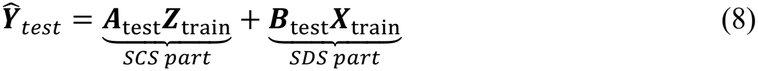

Where, 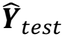 is the individual-level dFNC for the test group, and ***A***_*test*_, is the mixing matrix of the SCSs for the test group, learned from the train-group SLCs (***Z***_*train*_). Similarly, ***B***_*test*_ is the mixing matrix learned to decompose the dFNC into the SDSs (***X***_*train*_). We then determined the mean square error (MSE) for the reconstruction of ***Y***_*test*_, as well as for the SCS and SDS parts separately.

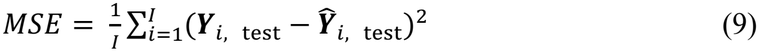

### Multimodal temporal dynamic parameter quantization

To characterize the temporal expression of the estimated states, i.e., the temporal dynamic parameters [15, 22, 33], we derived five dynamic measures from the state time courses (mixing matrices *A* and *B*): mean dwell time, frequency, occupancy rate, power, and variability. Because DECODE yields continuous loadings with arbitrary sign rather than discrete state assignments, state assignment was based on the magnitude of the loadings. At each timepoint, the state with the largest absolute loadings was considered occupied. Mean dwell time was defined as the average duration for which a state was occupied by a subject, frequency quantified the number of occurrences of a state over time, and occupancy rate represented the proportion of time points occupied by each state. Variability was estimated as the standard deviation of the state time course, and power as its mean squared amplitude. Mean dwell time, frequency, and occupancy rate summarize discrete state occupancy under a winner-take-all assignment. Variability and power, by contrast, are computed directly from the continuous loadings and therefore do not require discretization, characterizing graded state expression over time. Together, these measures summarize complementary aspects of temporal state expression, and allow the proposed decomposition to be evaluated not only spatially, but also temporally.

### State-wise group comparison and statistics

Group differences were evaluated across three pairwise diagnostic comparisons: cognitively normal (CN) vs mild cognitive impairment (MCI), MCI vs AD and CN vs AD. To consider the within-subject dependences, a linear mixed effect model (LMM), contorting for the confounds including age, sex, head motion, and site effects was applied to the cell-wise connectivity features and temporal dynamic parameters, followed by false discovery rate (FDR) correction on resulting P-values. Statistical analyses were performed separately for each structure-coupled and structure-decoupled state to assess differences in connectivity organization. We also examined group differences in the temporal dynamics of each state using the derived measures of dwell time, frequency, occupancy, power, and variability.

## Results

### Single hold-out validation and representational performance DECODE model

Figure 1 shows the main steps of the proposed DECODE model. We used diffusion MRI and functional MRI to identify ten recurring brain states that represent a structural-to-functional continuum of brain connectivity patterns, termed structure-coupled and structure-decoupled states. The proposed DECODE model was evaluated using a single hold-out pipeline, in which all samples were randomly divided into train and test groups. The test group included 30% of the subjects, and the remaining 70% comprised the train group.

Figure 2 (top row, left to right) shows the similarity (Pearson’s correlation) between the structural latent components (SLCs) estimated from the test group and those from the full sample, between the SLCs from the train group and those from the full sample, and between the SLCs from the test and train groups. The SLCs estimated from the test group (range: 0.65–0.93) and train group (range: 0.75–0.94) were highly similar to the corresponding SLCs from the full sample. Corresponding SLCs from the test and train groups were also correlated, with values ranging from 0.57 to 0.98. All structural patterns showed high correlation (r > 0.80), with the exception of SLC-1, which showed similarity below 0.80. This comparison of group-specific SLCs against those derived from the full sample confirms the generalizability of the estimated reproducible and stable SLC patterns.

**Figure 2.**
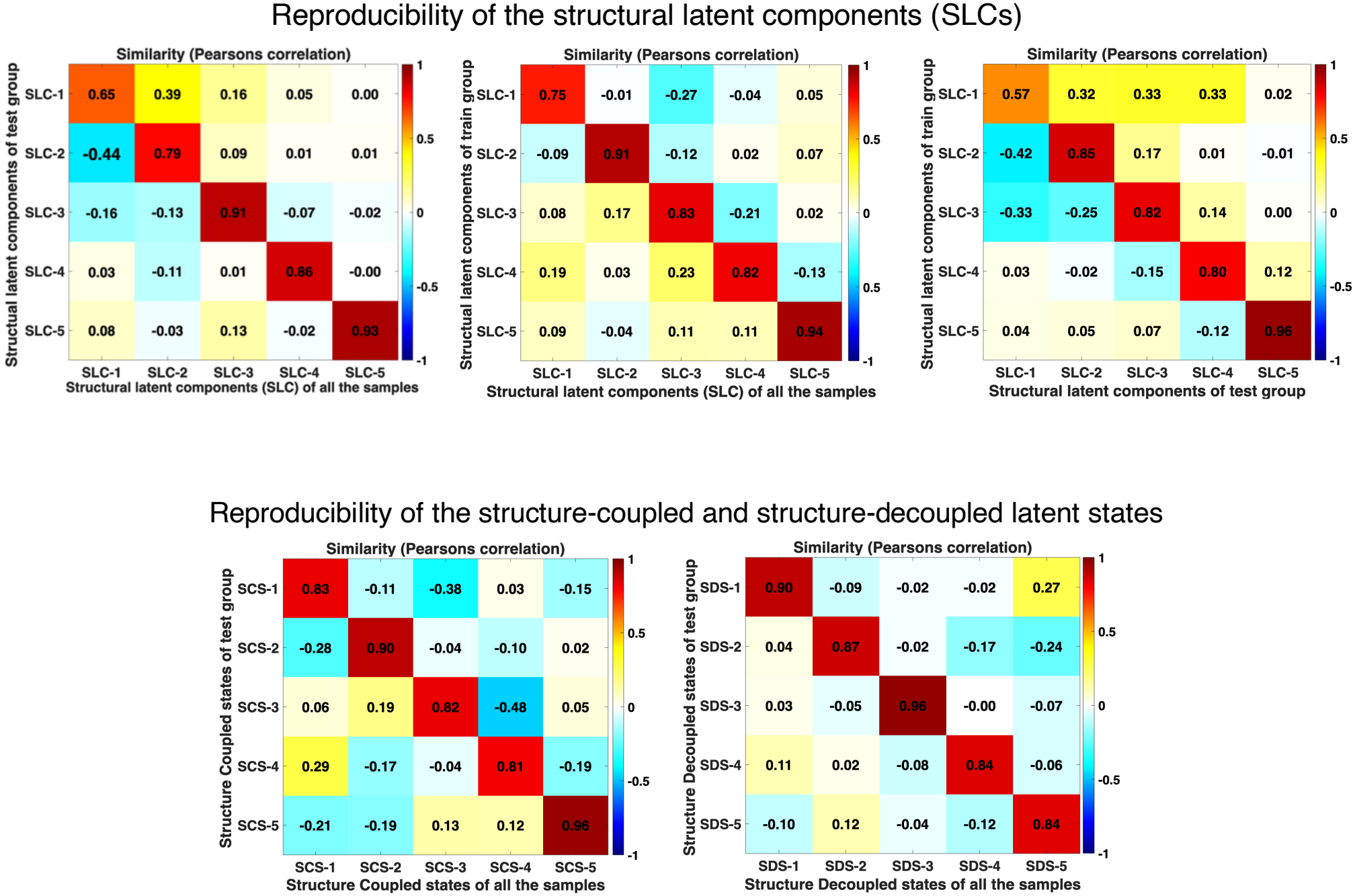
Model validation. Using a single hold-out approach, samples were split into train (70%) and test (30%) groups. **Top row:** similarity (Pearson’s correlation) of structural latent components (SLCs) between all samples and the test group, all samples and the train group, and the test group and the train group (left to right). **Bottom row:** similarity of structure-coupled (SCS) and structure-decoupled (SDS) states between the full sample and the test group, where test-group SCS/SDS were derived using train-group SLCs as priors. States were reordered so corresponding states align; high diagonal values indicate strong correspondence between test-group and full-sample results. Together, these results demonstrate the model’s generalizability to prior information from independent samples, along with their reproducibility and stability.

In Figure 2 (bottom row), we examined the renormalizabilty of the structure-coupled and structure-decoupled states. To this end, we computed the similarity between the structure-coupled and structure-decoupled states of the test group—decomposed using train-group SLCs—and the corresponding states from the full sample. The structure-coupled states of the test group, estimated using train-group SLCs, showed high correlation with the structure-coupled states from the full sample (range: 0.81–0.96 across corresponding states, Q_i_>0.8; left panel). Similarly, the structure-decoupled states estimated from the test group using train-group priors showed high correlation with the corresponding states from the full sample (range: 0.84–0.96, q_i_>0.8; right panel).

Furthermore, we computed the explained variance for the test and train groups using different prior information. For the inter-group analysis, using test-group SLCs to decompose the train-group dFNC and train-group SLCs to decompose the test-group dFNC, a substantial proportion of variance in each group’s dFNC was explained by the SCSs and SDSs (Figure 3, left panel). This analysis revealed lower explained variance for the SCSs (mean = 0.11 across test and train groups) than for the SDSs (mean = 0.30) in both the test (light-blue bars) and train (mid-blue bars) groups. Moreover, intra-group decomposition which each group’s dFNC was decomposed using its own SLCs, showed explained variance is consistent with the inter-group analysis, with only small variations (Figure 3, top-right panel). Compared with the SCSs and SDSs derived from the full sample (dark-blue bars), both the inter-group and intra-group analyses yielded similar levels of explained variance. We also compared the explained variance of the multimodal DECODE decomposition with that of a traditional unimodal functional-state decomposition, both computed on the full sample. The explained variance of the unimodal model (mean = 0.36) was slightly higher than that of either the SCSs (mean = 0.10, across test, train, and full-sample groups) or SDSs (mean = 0.30 across test, train, and full-sample groups) alone, but lower than their linear combination (mean = 0.40), consistent with the underlying assumption of the DECODE decomposition.

**Figure 3.**
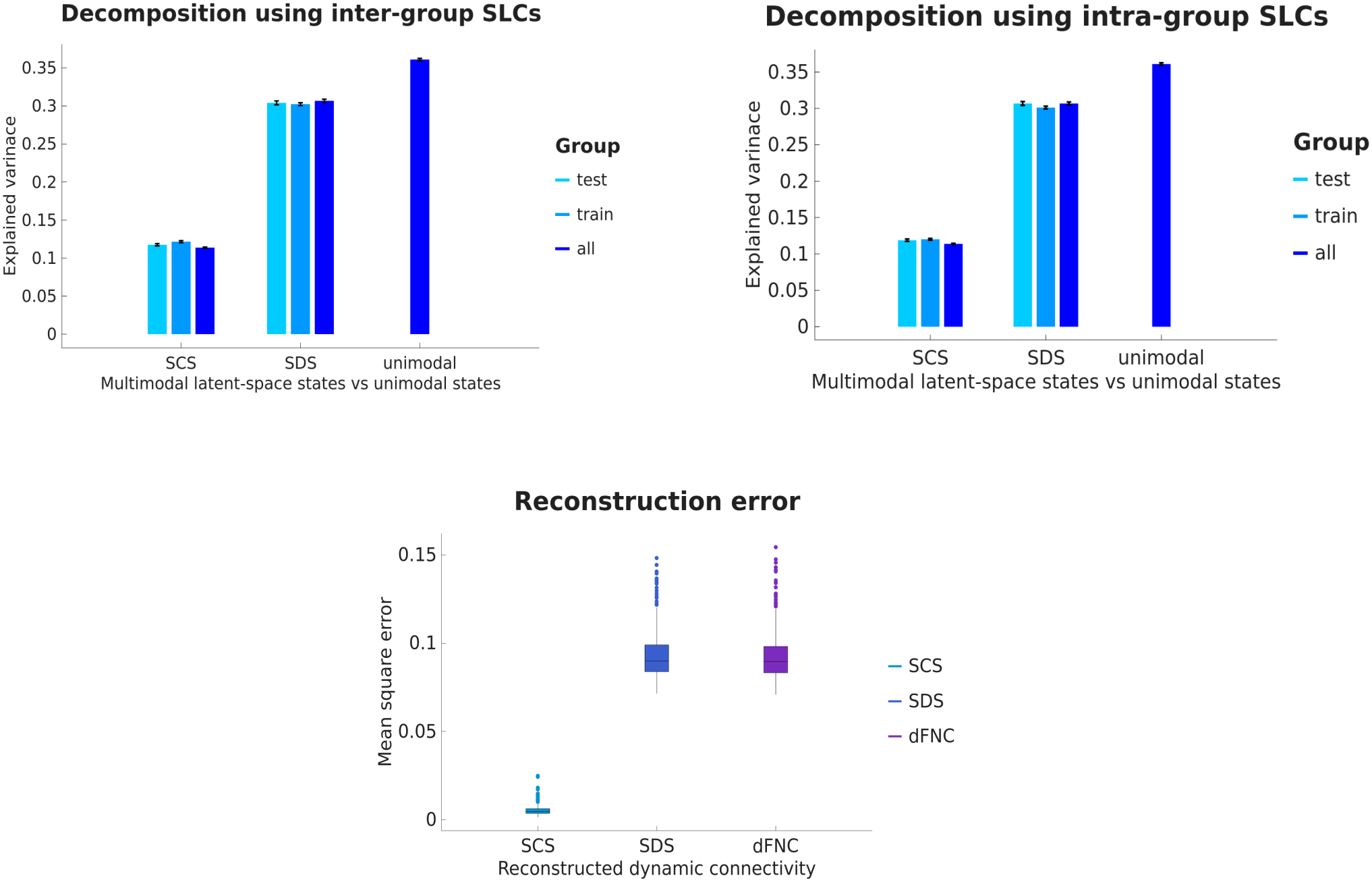
Model validation. Using a single-hold-out approach, samples were divided into train (70%) and test (30%) groups. **Top-left panel:** explained variance of the dFNC by SCS and SDS using inter-group SLCs — the test-group dFNC was decomposed using train-group SLCs, and the train-group dFNC was decomposed using test-group SLCs. Bars show mean explained variance across time for the test group (light blue), train group (mid-blue), and full sample (dark blue), with standard error (black lines); a unimodal decomposition (dFNC only) is shown for comparison. The multimodal model (**mean SCS** = 0.11 + **mean SDS** = 0.30) explained more variance than the unimodal model (mean = 0.36). **Top-right panel:** explained variance using intra-group SLCs — each group’s dFNC was decomposed using its own group’s SLCs — yielding values nearly identical to the inter-group results. **Bottom panel:** reconstruction error (MSE) for the test group using train-group SLCs, shown for SCS alone (bluish green; **mean** = 0.0047, 25th–75th percentile: 0.0036–0.006), SDS alone (blue; **mean** = 0.089, 25th–75th percentile: 0.083–0.1), and combined SCS+SDS (purple; **mean** = 0.089, 25th–75th percentile: 0.083–0.98). All MSEs were below 0.1, indicating reasonable reconstruction accuracy for the proposed DECODE model.

In addition, we estimated the reconstruction error for the test-group decomposition using train-group SLCs, displayed as boxplots in Figure 3 (bottom panel). Reconstruction error, quantified using the mean square error (MSE), was computed separately for the SCS component, the SDS component, and the full dFNC reconstruction. The bluish-green boxplot represents the MSE for SCS reconstruction (mean = 0.0047; 25th–75th percentile: 0.0036–0.006), the blue boxplot represents the MSE for SDS reconstruction (mean = 0.089; 25th–75th percentile: 0.083–0.1), and the purple boxplot represents the MSE for the combined SCS+SDS reconstruction (mean = 0.089; 25th–75th percentile: 0.083-0.098). These results indicate reasonably low MSEs (<0.1) for reconstructing functional dynamic connectivity using the proposed DECODE model. While the SCSs showed the lowest reconstruction error, the majority of the total error was attributable to the SDSs. In sum, the explained variance and MSE results collectively demonstrate the efficiency of the proposed DECODE model in decomposing brain dynamic connectivity into meaningful states.

### DECODE reveals temporal continuum structure-coupled and structure-decoupled functional latent-space states

Figure 4 illustrates the structural priors and the ten recurring brain latent-space states, transition from structure-coupled to structure-decoupled. From left to right shows the transition from structural latent-space components (SLCs) to structure-coupled states and to structure-decoupled states.

**Figure 4.**
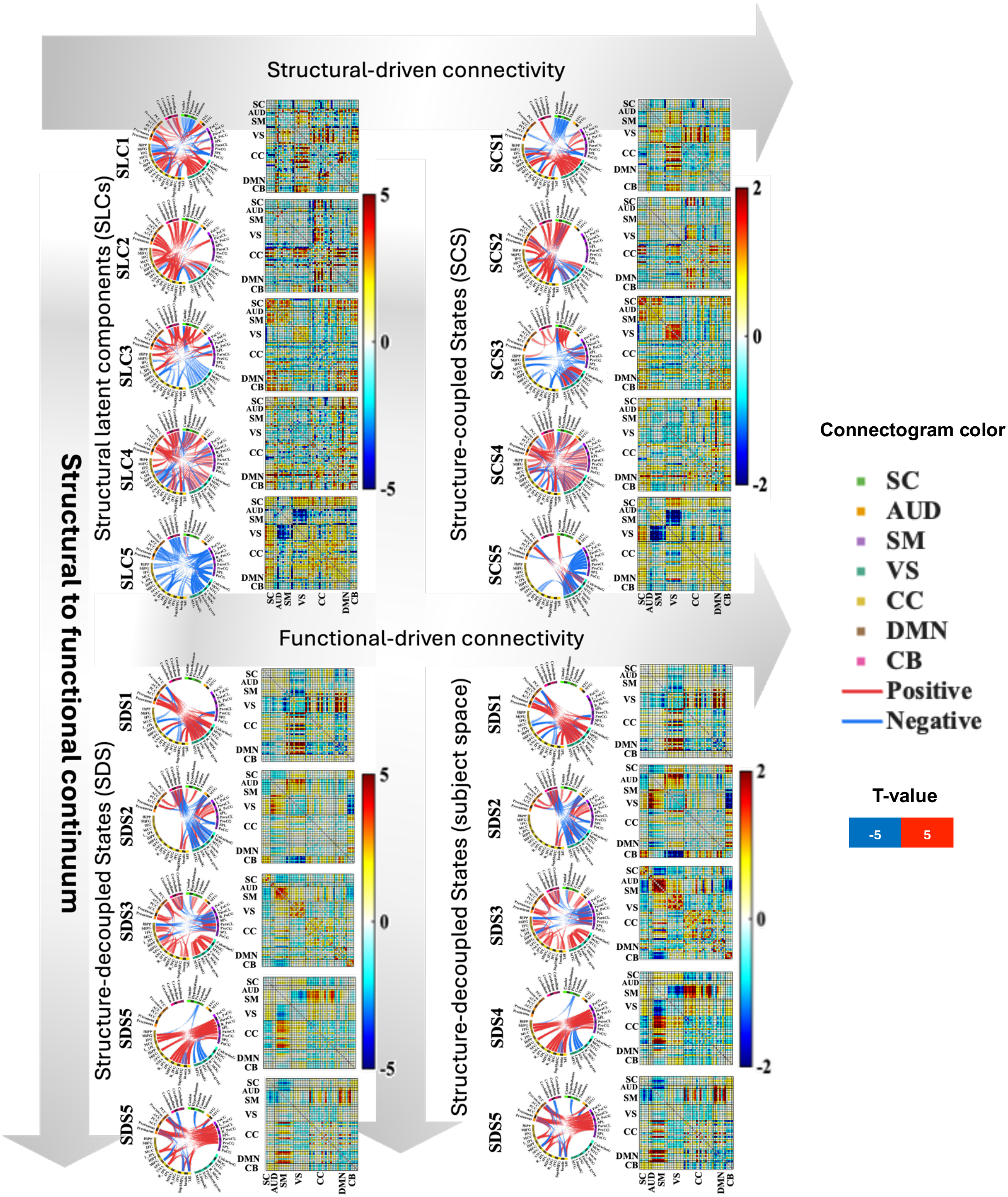
Temporal-continuum of structure-coupled and structure-decoupled functional latent-space states. Structural connectivity latent space components (SLCs) (left panel) represent dominant structural connectivity patterns, derived from the structural connectome. Incorporating these priors into the model yields multimodal states (middle panel) that remain aligned with the structural architecture (structure-coupled states, SCS). After reconstructing unexplained functional information from structural using regression analysis, the resulting patterns reveal structure-decoupled functional states, SDS. (right panel), reflecting interactions that are progressively less constrained by structural information. The arrows indicate progression along from the structural information to the well coupled structural-functional information (structure-coupled states) to the less coupled structural-functional information (structure-decoupled states). Connectograms display the top 5% strongest connections (red: positive; blue: negative), with chord width representing connection strength. The adjacent matrices show the full connectivity structure.

Each of the five structural priors, SLCs stands for different covarying patterns; SLC1: VS and AUD domain covariation with other domains. SLC2: CC covariation with other domains. SLC3: sensory and SC with DMN and CB. SLC4: DMN, CB and SC. SLC5: SC, intra-CC and VS-SM. These observations suggest that the estimated functional states inherit topological features from the structural priors, supporting the notion that structural connectivity provides an anatomical scaffold that constrains large-scale functional brain dynamics.

By incorporating the structural information into the model, the estimated structure-coupled functional states (second column on top) reflect connectivity configurations that remain strongly aligned with the underlying structural architecture, representing structure-coupled functional states. Structural connectivity bases and the structure-coupled functional states reveal substantial similarity in their network organization. Several large-scale connectivity motifs, including long-range interactions linking frontal, parietal, and temporal regions, are preserved across the two representations. In addition, key hub regions such as the precuneus/posterior cingulate and inferior parietal cortex exhibit high connectivity in both structural and functional patterns.

As the structurally explained information is modeled, the unreconstructed connectivity patterns give rise to structure-decoupled functional states, which capture functional interactions that are not explained by the direct structural connectivity. Compared with the structure-coupled functional states, the structure-decoupled states exhibit reduced similarity to the structural connectivity latent space. While several connectivity motifs observed in the structural latent space are preserved in the structure-coupled states, in structure-decoupled states, connectivity patterns are diverged from the priors. As, SDS1: VS, SDS2: VS-SM and CB, SDS3: intra-domain, SDS4: SM-VS and SM-CC. SDS5: SM. The decoupled states show more heterogeneous and selective connectivity configurations, and the emergence of novel interaction patterns not directly reflected in the structural latent components. These findings suggest that functional brain dynamics can be transformed to a two set of complimentary latent space, a space well coupled with structural information and a space less coupled. The connectogram in Figure 4 illustrate the top five percent strongest connections within each state, while the corresponding connectivity matrices provide a complete representation of the network structure.

In addition, we determined the amount of dFNCs information that is explained by brain states; from structure-coupled to structure-decoupled. To this end, a window-wise, explained variance (R^2^) was computed by fitting each window to the five structural bases, five structure-coupled and five structure-decoupled states through a linear regression model. While the structure-decoupled has the highest explained information (Mean ± standard error: 0.33± 0.01), the structure coupled states, explained less than the structure-decoupled states (Mean ± standard error: 0.15± 0.01). Results align with findings in Figure 3, as the amount of time-resolved dynamic functional information explained by the states increases from a structure-coupled to structure-decoupled functional states. Overall, results show an increasing consistency from structural information to well aligned structure-function (structure-coupled) to less aligned structure-function (structure-decoupled) information, achieved by decreasing the structural information and increasing the functional information.

### DECODE reveals Alzheimer-related pathology increasingly affects brain functional dynamic states beyond what can be explained by functional alone

State-wise statistical comparisons of structure-coupled and structure-decoupled connectivity states using linear mixed effect model (LMM) are shown in Figure 5 and Figure 6. Each row corresponds to a state, and each column indicates the three diagnostic contrasts, CN-MCI, MCI-AD, and CN-AD that has been investigated in this work. Connectogram display the statistically significant connections (FDR-corrected P-values<0.05), with red and blue chords indicating positive and negative *T-values*, respectively, and chord width reflecting magnitude of the *T-values*.

**Figure 5.**
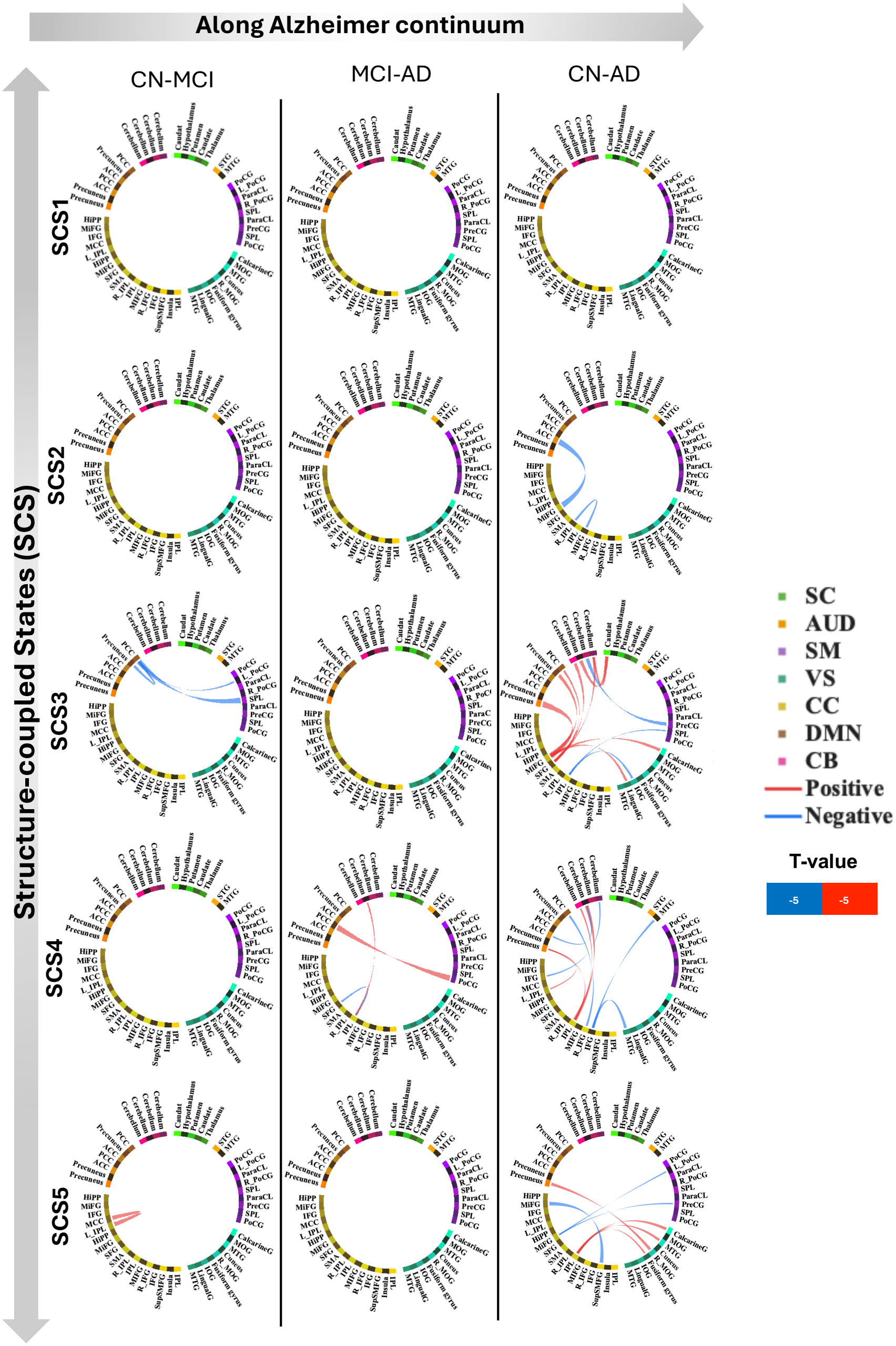
State-wise statistical comparison of structure-coupled functional latent-space states across the Alzheimer disease progression. Rows correspond to individual states, and columns represent the three diagnostic contrasts, CN-MCI, MCI-AD, and CN-AD, shown for linear mixed effect model (LMM). Statistical testing was performed in a cell-wise manner for each connectivity feature, and only significant connections (FDR-corrected P-value < 0.05) are displayed. Connectograms depict the significant connections, with red and blue chords indicating positive and negative T-values, respectively, and chord width reflecting magnitude of the T-value. Overall, group differences were state-dependent and increased with disease severity, with the most widespread alterations observed in CN-AD, intermediate effects in MCI-AD, and fewer differences in CN-MCI.

**Figure 6.**
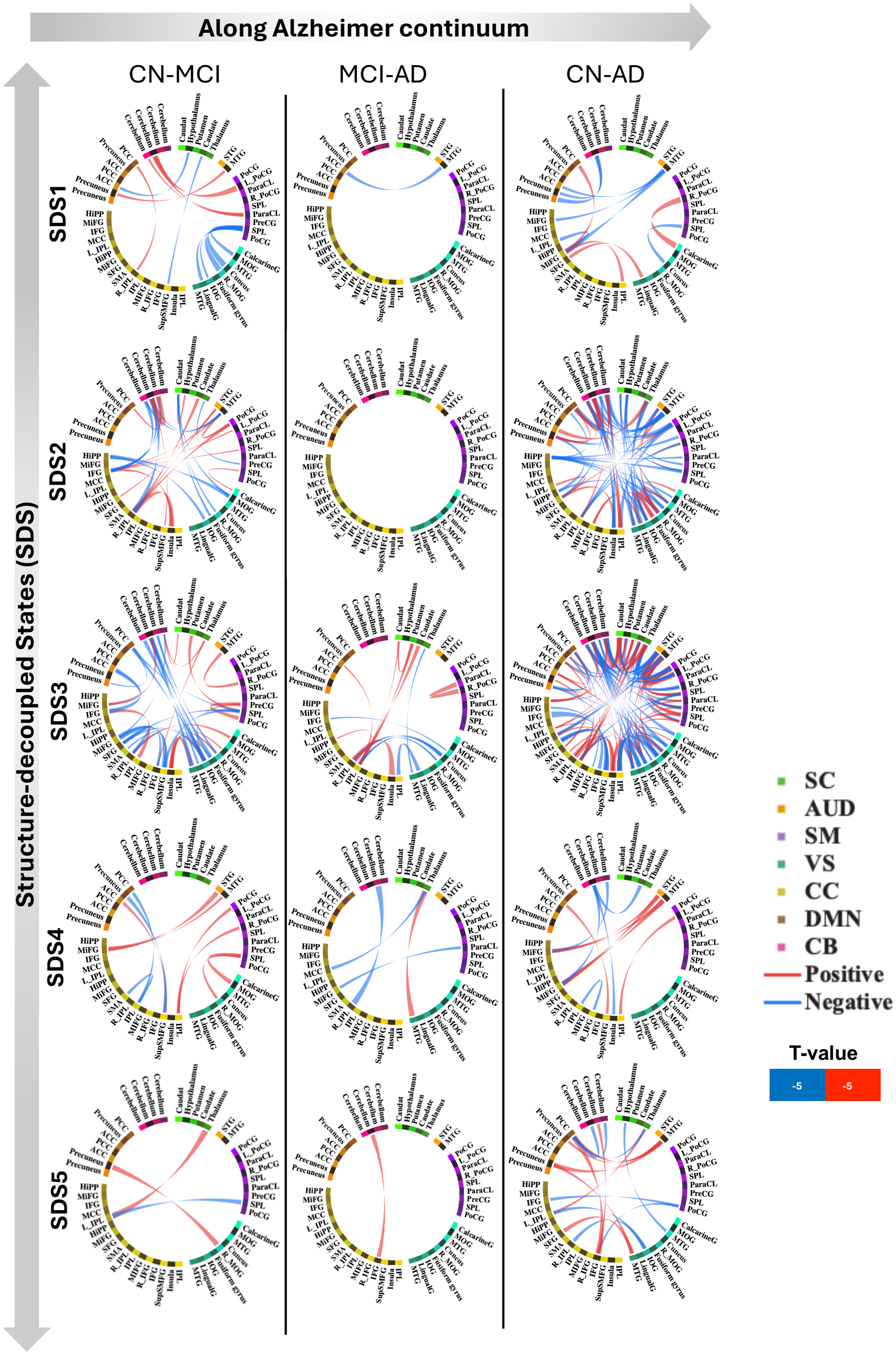
State-wise statistical comparison of structure-decoupled functional latent-space states across the Alzheimer disease progression. Rows correspond to individual states, and columns represent the three diagnostic contrasts, CN-MCI, MCI-AD, and CN-AD, shown for linear mixed effect model (LMM). Statistical testing was performed in a cell-wise manner for each connectivity feature, and only significant connections (FDR-corrected P-value < 0.05) are displayed. Connectograms depict the significant connections, with red and blue chords indicating positive and negative T-values, respectively, and chord width reflecting magnitude of the T-value. Similar to the structure-coupled, structure-decoupled group differences were also state-dependent and increased with disease severity, with the most widespread alterations observed in CN-AD, intermediate effects in CN-MCI, and fewer differences in MCI-AD. However, in comparison, the structure-decoupled states exhibited stronger abnormalities than the structure-coupled states, indicating that disease progression is associated with increasingly disrupted functional configurations that are less constrained by the structural scaffold.

Results reveal a state-dependent pattern of altered connectivity across the Alzheimer’s progression. The CN-AD comparison showed the most widespread and strongest differences, MCI-AD showed intermediate alterations, and CN-MCI showed the fewest, consistent with progressively increasing disruption from normal cognition to dementia. These effects were stronger in a subset of states rather than uniformly distributed, indicating selective vulnerability of specific dynamic configurations.

Notably, the strongest and most extensive group differences were concentrated in the structure-decoupled states (Figure 6), whereas the structure-coupled states (Figure 5) showed relatively sparse and localized abnormalities. Among the structure-coupled states, SCS3 (structural modularity in SC, CB and sensory domains based on Figure 4) was most affected in CN-MCI, whereas SCS3 and 4 (structural modularity in SC, DMN and CB domains based on Figure 4) showed stronger involvement in MCI-AD. In contrast, the structure-decoupled states exhibited broader abnormalities overall, with SDS1, 2, 3, and 4 (functionally modular in VS, DMN, CB, sensory and CC based on Figure 4) involved in CN-MCI and SDS2 and 3 particularly affected in MCI-AD. The CN-AD comparison showed significant differences across nearly all coupled and decoupled states, with both increased and decreased connectivity patterns, indicating substantial disruption of state-specific network organization in dementia. Interestingly, the abnormalities (significantly different between groups) are not purely DMN-centered, however involvement of VS, AUD, SM, CC, DMN and CB may suggest a broader reorganization than the usual reported posterior DMN disruption.

Statistical test results at the domain level, show that the most recurrent abnormalities involved sensorimotor, visual, cerebellum, and subcortical systems, with additional involvement of DMN and cognitive control interactions. Findings support an organized progression of dynamic connectivity disruption in Alzheimer’s disease, shifting from sparse, state-specific abnormalities in earlier stages to broader cross-network dysconnectivity in dementia.

### Temporal dynamic parameter quantification shows AD-related temporal dysregulation

Quantification of the temporal dynamic activity of each state through the five metrices including Mean dwell time, frequency, occupancy rate, power and variability by using the state-specific occurrence (activity) over time is illustrated in Figure 7. Results demonstrated a distributed significant difference (FDR-corrected P-value<0.05) across both structure-coupled (SCS) and structure-decoupled (SDS) states as of the statistical analysis and are distinguished by a bracket and star in Figure 7. While dynamic parameter of the structure-coupled states mostly shows significant disconnection (FDR-corrected P-value<0.05) between the CN vs AD groups for the frequency, occupancy, power and variability, the structural-decoupled states reflect differences for the CN vs MCI, CN vs AD and CN vs MCI over dynamic parameters.

**Figure 7.**
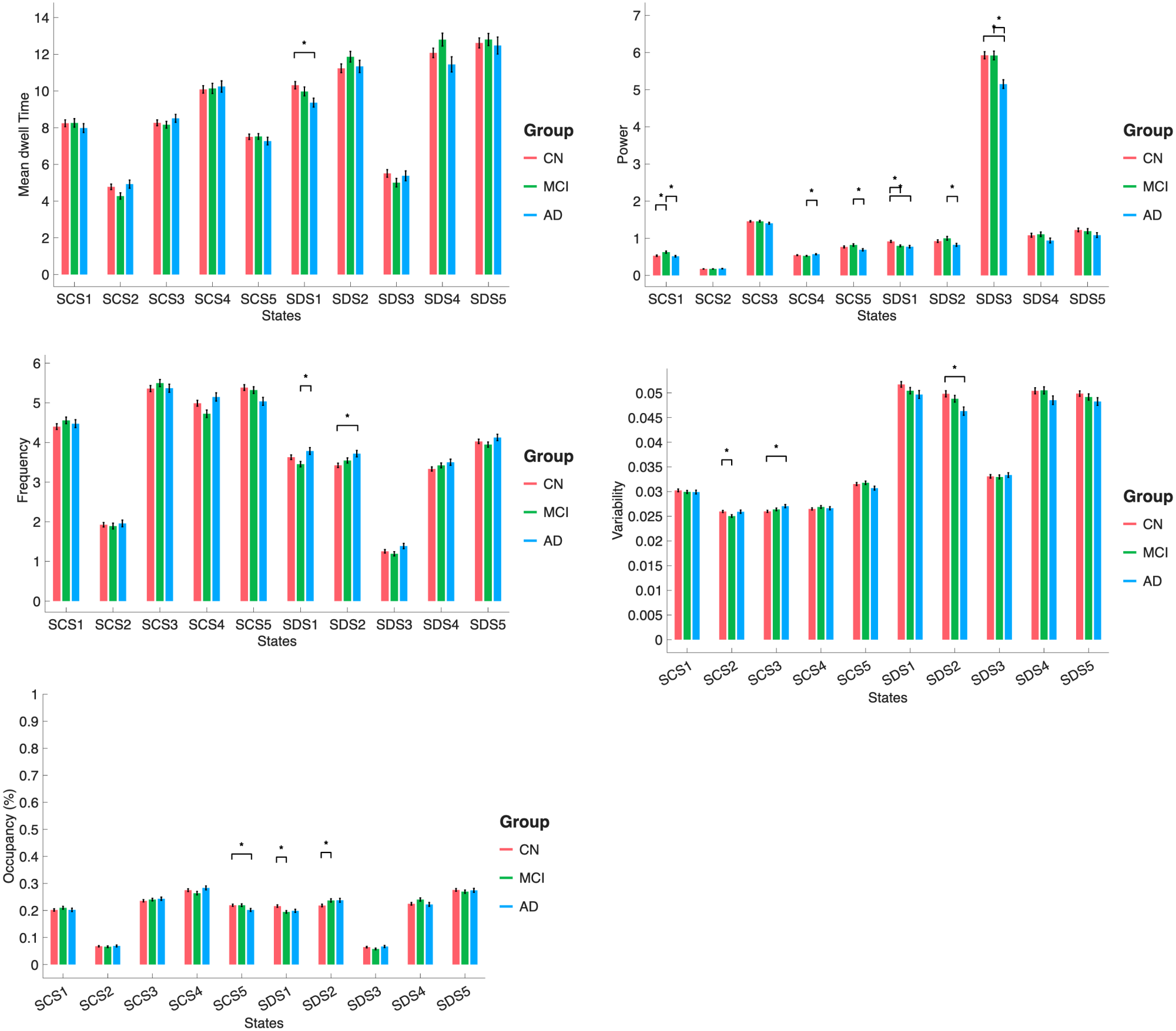
Quantification of temporal dynamic parameters. We quantified the variation in activity profile for the ten structural coupled (SCS) and decoupled (SDS) states over time– called temporal dynamic parameters– through five metrices, Mean dwell time, Frequency, Occupancy rate, Power and variability of the time courses at five panels. Quantification analysis followed by statistical test determined significant variation between three groups of subjects (control: CN, red bars, mild cognitive impairment: MCI, green bars and dementia: AD, blue bars) according to estimated parameters. Significant (FDR-corrected P-value<0.05) states for each of the disease-pair are distinguished by a bracket and star. Overall, there are significant differences between groups regarding each temporal dynamic parameter that mostly are observed in the Frequency, Occupancy rate, Power and Variability.

Among the states, SCS5, SDS1 and SDS2 are the most contributed ones, dominantly related to connectivity patterns for the visual domain with subcortical, sensory motor and cerebellum– VS-SC, VS-SM, VS-CB– and sensory motor connection with cognitive control (SM-CC). Interestingly, it is observed that DMN related states (SCS3, SCS4 and SDS2) mostly affected the Frequency, Variability and Power.

In Figure 7, top left, Mean dwell time with a significant difference for the CN vs AD in the structure-decoupled states 1, is the minimum significancy among all the parameters. In the mean dwell time panel, significant effects are visible mainly in the decoupled portion, indicating altered persistence of these states across groups. Interestingly, frequency showed significant differences in structure-coupled and structure-decoupled states for the CN vs AD and MCI vs AD. In the frequency and occupancy panels, several significant differences are observed around SDS1-SDS2, consistent with diagnosis-related shifts in how often these states are visited and how much time is spent in them. On the other hand, power–Figure 7, top right, shows the highest number of state-level significance for all three-diagnosis pairs among all the parameters. The power of time courses shows the clearest single effect, with a striking peak in SDS3 and a significant group difference, indicating that this state is especially sensitive to disease-related change. By comparison, variability of the time courses shows fewer and smaller between-group effects, although significant differences are still present in selected states, suggesting that the disease signal is stronger in state engagement and persistence than in overall temporal dispersion. The most prominent abnormalities are concentrated in the structure-decoupled states, particularly SDS1-SDS3, suggesting that dementia progression is associated not only with altered state expression, but also with disturbed temporal regulation of states that are less directly constrained by the structural.

## Discussion

In the present study, we introduced DECODE, a novel multimodal data-driven model designed to move beyond conventional approaches for studying structure-function relationships using static coupling measures, post hoc multimodal comparisons, or functional-only state estimation. The central methodological contribution of DECODE is its ability to decompose dynamic functional brain organization into structure-coupled and structure-decoupled recurring states, thereby positioning recurrent brain dynamics along a temporal-continuum structural coupling. Rather than reducing the structure-function relationship to a single coupling score, DECODE provides a state-wise decomposition that explicitly separates functional patterns that remain aligned with the anatomical scaffold from those that are relatively less constrained by it. In this way, the framework offers a more mechanistic account of how dynamic brain organization reflects the interplay between structural constraint and functional flexibility, and it is not restricted to a single dataset or disease context [34-36].

### DECODE Reliably Captures Generalizable and Reproducible structure–function network dynamics across temporal continuum

The single-hold-out validation supports the generalizability and reproducibility of the proposed model. SLCs estimated separately from the test and train subsamples closely matched those derived from the full sample (r = 0.65–0.94), and this held across nearly all components (r > 0.80, except SLC-1), indicating that the structural patterns are stable rather than sample-specific. Structure-coupled and structure-decoupled states, temporal-continuum connectivity patterns occurred repeatedly over time, estimated using inter-group priors, meaning test-group states derived from train-group SLCs, also correlated highly with the full-sample results (r = 0.81–0.96). This suggests the SLCs are not overfit to the sample they were estimated from, and that the functional-structural coupling patterns they capture generalize to unseen data.

The explained-variance and mean square error results add further support. Explained variance was similar across the inter-group and intra-group SLCs used for decomposition, highlighting the model’s independence from the specific sample or sample size used to estimate the structural priors. This has practical implications: structural priors derived from one sample could plausibly be applied to functional data from another without a meaningful drop in performance, without requiring matched structural and functional data from the same individuals. The combined SCS and SDS also explained more variance than either alone, and modestly more than a unimodal functional decomposition, consistent with the model’s premise that structure-coupled and structure-decoupled dynamics carry complementary information. Finally, reconstruction error for the test group using train-group parameters remained low (MSE < 0.1), though most of this error came from the SDS rather than the SCS, suggesting that structure-decoupled dynamics may be less structurally constrained and worth examining further in future work. Overall, these results support the model as a reproducible and generalizable approach to characterizing structure-function coupling in brain dynamics, providing a validated basis for using the SLCs, SCS, and SDS estimated from the full sample in the subsequent statistical analyses.

### DECODE reveals temporal continuum structure-coupled and structure-decoupled functional latent-space states

Figure 4 provides the additional conceptual validation of this framework by showing that the estimated states are distributed along a graded temporal-continuum structure-coupled and structure-decoupled rather than falling into a binary division between structurally determined and functionally independent processes. The structure-coupled states retained major topological motifs of the structural bases, including large-scale patterns consistent with the underlying anatomical organization, indicating that structural connectivity constrains important features of recurrent functional brain states. By contrast, the structure-decoupled states displayed more selective, heterogeneous, and functionally driven connectivity configurations, suggesting that a substantial portion of brain dynamics emerges beyond direct structural alignment. This pattern is consistent with prior work showing that anatomical connectivity constrains, but does not fully determine, large-scale functional organization [37], and that this relationship varies systematically along cortical gradients and hierarchies [7, 13, 38, 39]. It also aligns conceptually with recent dynamic multimodal efforts that seek to derive structural bases that adapt to functional organization rather than treating structure only as a static reference [34].

Estimation of the explained variance complements this observation by quantifying how much functional information is captured across the proposed temporal continuum. The structure-coupled states explained 0.11± 0.01 (Mean ± standard error) of the dFNCs, and the structure-decoupled states explained the largest proportion (Mean ± standard error: 0.32± 0.01). This progressive increase in explained functional variance supports the interpretation that structural representations provide an important anatomical backbone, but that more functionally expressed and less structurally constrained states capture a larger share of transient functional organization. Importantly, the structure-coupled states occupied an intermediate position, preserving measurable structural information while also incorporating dynamic functional variability. Taken together, findings support the view that brain dynamics are neither fully fixed by structure nor entirely independent of it, but instead unfold along a graded temporal continuum from structural constraint to functional flexibility [13, 36, 40-42].

This temporal continuum framework offers a principled way to distinguish between anatomically anchored and relatively decoupled components of dynamic functional organization. Such a distinction is relevant not only for neurodegenerative disease, but also for broader questions in systems neuroscience concerning how structural architecture shapes, but does not fully define, transient brain dynamics. In contrast to standard structure-function coupling approaches, which typically summarize the relationship between modalities using regional or static descriptors, DECODE provides a dynamic and state-wise representation of that relationship. This makes the framework potentially useful for investigating interindividual variability, developmental change, cognitive flexibility, and multimodal dysfunction across a wider range of neurological and psychiatric conditions [7, 36, 39].

### DECODE reveals Alzheimer-related pathology increasingly affects brain functional dynamic states beyond what can be explained by functional alone

Within this broader framework, the application to Alzheimer’s disease serves as an informative validation setting. Our results showed that the most pronounced group differences in connectivity organization were concentrated in the structure-decoupled states, whereas structure-coupled states exhibited relatively fewer abnormalities. This pattern suggests that disease-related effects may become most visible in transient functional configurations that are less tightly constrained by the structural scaffold. Our findings– as of the Figure 5 and Figure 6– are consistent with prior evidence that Alzheimer’s disease is characterized by progressive disruption of large-scale functional organization, particularly involving the default mode and executive control systems, together with abnormalities in dynamic connectivity and altered structure-function relationships. Previous work [43] has shown that AD and MCI are associated with reduced flexibility of dynamic functional network configurations, altered occupancy of specific connectivity states, and impaired interactions among visual, sensorimotor, and auditory systems, supporting the view that AD is a disconnection syndrome extending beyond the DMN alone [43]. In parallel, recent multimodal studies have reported abnormal structure-function coupling across the AD progression, indicating that the correspondence between anatomical wiring and functional coordination becomes progressively disturbed in disease [44]. Against this background, our results extend previous findings by showing that the most pronounced group differences were concentrated in the structure-decoupled dynamic states, whereas structure-coupled states exhibited relatively sparse abnormalities. This suggests that disease progression may increasingly affect transient functional coordination that is less constrained by the structural scaffold. The observed involvement of sensorimotor, visual, cerebellar, subcortical, DMN, and cognitive control domains is also in line with prior reports of cerebellar connectivity disruption in AD/MCI and broader cross-network reorganization beyond canonical memory-related systems [45]. Together, these results support the potential of the model in interpretation of the underlying dynamic interactions and uncovering hidden patterns changed across brain disorders such as the AD. Specifically, this study showed that AD progression reflects not only weakening of established network architecture, but also increasing dysregulation of flexible, state-dependent interactions across sensory, associative, and cerebellar systems [46].

### Temporal dynamic parameter quantification shows AD-related temporal dysregulation

The temporal dynamic analyses further support the usefulness of the proposed decomposition. Findings in Figure 7, suggest that dementia-related abnormalities in brain dynamics are not distributed uniformly across states but instead are concentrated primarily in the structure-decoupled states, especially SDS1-SDS3. This pattern indicates that disease progression may involve more than a change in the strength of state expression alone; it also appears to affect the temporal organization of brain states that are less tightly constrained by the underlying structural scaffold. In other words, as individuals progress from CN to MCI and dementia, the disturbance seems to be expressed most strongly in the timing, persistence, and recurrence of functional configurations that are relatively freer from direct anatomical constraint. This interpretation is broadly consistent with prior work showing that Alzheimer’s disease is characterized by abnormalities in dynamic functional connectivity, including altered dwell time, occupancy, recurrence, and state transitions, rather than by static connectivity disruption alone. Studies of recurrent functional connectivity states in AD have shown that patients can spend less time in some states and more time in others, supporting the idea that neurodegeneration produces a redistribution of state engagement rather than a simple monotonic increase or decrease in dynamical measures [47]. More generally, this result demonstrates that DECODE is informative not only at the level of spatial state organization, but also at the level of temporal expression, further supporting its value as a general framework for multimodal dynamic brain-state analysis [7, 34, 36, 39].

### Limitation

This study provides a broader perspective on brain structure-function interactions, but several limitations should be acknowledged. Current implementation assumes a largely linear separation between structure-coupled and structure-decoupled functional patterns, whereas the true relationship between brain structure and function is likely more complex and potentially nonlinear. As a result, the identified states and their interpretation may still be influenced by the selected node definition and spatial scale. Future work may therefore benefit from nonlinear decomposition strategies. Such extensions would further strengthen DECODE as a generalizable framework for studying how dynamic brain function is constrained by, and diverges from, anatomical structure.

## Conclusion

In summary, we introduced DECODE, a data-driven latent-space framework that decomposes dynamic functional connectivity into temporally recurring structure-coupled and structure-decoupled brain states. By moving beyond regional structure-function coupling measures, DECODE provides a state-wise framework for characterizing how brain dynamics are constrained by, and diverge from, the structural connectome. Applied to Alzheimer’s disease, we showed that structure-coupled states preserved major structural motifs, whereas structure-decoupled states captured more heterogeneous functional patterns, explained greater functional variance, and were more sensitive to disease-related abnormalities. These findings support DECODE as an efficient and generalizable framework for identifying recurring structure-dynamic connectivity patterns beyond a single dataset or brain disorder.

## Code Availability

The main codes are already available within GIFT toolbox (https://github.com/trendscenter/gift), and the unified framework will also be released as a module within GIFT.

## Acknowledgment

This work was supported in part by the National Science Foundation (NSF), and by the National Institutes of Health (NIH).

